# Liver-Derived, Circulating Xanthine Oxidoreductase Drives Vascular Impairment Associated with Inhalation of Ultrafine Particulates

**DOI:** 10.1101/2025.05.07.652261

**Authors:** Xena M. Williams, Alec T. Bossert, Maddison Seman, Sara E. Lewis, Hailey McCallen, Josilyn Pilkerton, Elizabeth C. Bowdridge, Mark A. Paternostro, W. Travis Goldsmith, Timothy R. Nurkiewicz, Salik Hussain, Eric E. Kelley, Evan DeVallance

## Abstract

Inhalation of ultrafine particles (UFP) mediates systemic vascular impairment which is, in part, driven by elevated rates of oxidant generation. One significant source of oxidant production in the vascular compartment is the purine catabolizing enzyme, xanthine oxidoreductase (XOR). However, mechanisms linking XOR and/or endothelial glycosaminoglycan (GAG)-sequestered XOR to vessel dysfunction allied to UFP inhalation remain underexplored. Based on known interactions between UFP and the liver, we hypothesized that exposure could lead to hepatic release of XOR to the circulation which subsequently contributes to vascular impairment. Utilizing our murine hepatocyte-specific XOR knockout (XOR^Hep-/-^) model (loss of function) in conjunction with reintroducing exogenous XOR (restoration of function) we demonstrate a specific role for liver-derived XOR in the pathogenesis of UFP-induced vascular impairment. Exposure of mice as well as *in vitro* exposure of hepatocytes to our model UFP, nano titanium dioxide (nTiO_2_) results in the upregulation and active release of XOR. Drinking water supplemented with the XOR inhibitor febuxostat or nitrite (NaNO_2_^-^) partially prevented nTiO_2_-induced impairment of vascular reactivity. Interestingly, nitrite appears to cause a down-regulation of hepatic XOR. XOR^Hep-/-^ mice were partially protected against both impairment of endothelial dependent dilation and augmented angiotensin II constriction. To further demonstrate the role of circulating XOR in nTiO_2_-induced impairment of vessel reactivity, XOR^Hep-/-^ mice had circulating XOR restored by *i.v.* injection prior to exposure, which eliminated the protection of the hepatic knockout. It is important to note that acute restoration of intraluminal XOR in isolated vessels did not alter endothelial-dependent dilation or angiotensin II constriction. As such, we interrogated potential downstream mediators of XOR effects on endothelial function and found a decrease in the repressive trimethylation of lysine 9 on histone 3. Together these findings demonstrate that circulating XOR is a key contributor to endothelial dysfunction caused by UFP exposure. However, the impairment is not acute in nature and might involve epigenetic-mediated alterations in gene expression.

## Introduction

Airborne ultrafine particles (UFP; aerodynamic diameter <∼100nm) from both natural and man-made sources are increasing in populated areas worlwide^1^. This constitutes a considerable global healthcare issue as data affirms a relationship between UFP exposure and cardiovascular/ all cause morbidity and mortality. Growing evidence suggests UFP inhalation elicits chronic systemic low-grade inflammation similar to other disease processes^2^. Our group as well as others have experimentally demonstrated that UFP negatively impacts cardiovascular function^3–14^. These studies have implicated over-production of oxidants as contributory; however, the source of these oxidants remains poorly defined^6,10^.

Xanthine oxidoreductase (XOR) is an essential molybdoflavin enzyme in the purine catabolism pathway. XOR exists in either the oxidase (XO) or dehydrogenase (XDH) form, derived from a single gene (XDH). Originally discovered over a century ago^15–17^, the enzyme is converted from XDH to XO^18,19^, which utilizes O_2_ as an electron acceptor, generating oxidants (hydrogen peroxide (H_2_O_2_) and superoxide (O_2_^●-^)). Due to its proclivity to produce oxidants, the XO form has long been implicated in numerous disease pathologies. This was first appreciated in seminal studies of ischemia reperfusion injury^20–27^ resulting in XOR being identified as a contributor in numerous cardiovascular pathologies^28–34^. While XOR is expressed in the vascular endothelial cells, it is also free in circulation, where levels are known to increase in chronic and acute inflammatory conditions^23,24,35,36^. The prevailing dogma suggests that circulating XO binds to endothelial glycosaminoglycans (GAGs), producing O_2_^●-^ which subsequently diminishes nitric oxide (^●^NO) via rapid and direct reaction (O_2_^●-^ + ^●^NO ® O=NOO^-^). While widely accepted, very little durable experimental evidence supports this assumption whereas localization of O_2_^●-^ production (luminal face of endothelium) and biochemical properties (e.g. negative charge) question the feasibility. For example, it is not likely that negatively charged O_2_^●-^ can traverse the endothelial cell membrane and affect ^●^NO diffusion to the basolateral surface on its journey to the smooth muscle; however, alternative mechanisms linking XOR to vessel dysfunction have yet to be elucidated.

We recently demonstrated that the primary source of XO in the circulation is the liver where hepatocytes can be solicited to actively export XOR^37,38^. Interestingly, reports indicate UFP can cross into the circulation where they accumulate in the liver^39,40^ which poses the question, might UFP inhalation exposure stimulate the release of XOR from hepatocytes? Indeed, previous links have been made between toxicant exposures and XOR^41–43^; however, it remains unknown if UFP triggers hepatic release of XOR into the circulation. In addition, it is also unknown if XO binding to endothelial GAGs contributes to UFP-induced vascular impairment. As such, we examine herein if nano-titanium dioxide (nTiO_2_, our model UFP) induces hepatic release of XOR which then contributes to vascular impairment.

Utilizing both *in vitro* and *in vivo* approaches along with pharmaceutical inhibition, exploitation of alternative substrate-driven enzyme activity (nitrite reductase) and genetic deletion, we provide compelling evidence to suggest that UFP-driven vascular impairment is mediated by XO, which is prevented/restored by febuxostat treatment or nitrite supplementation. In addition, we provide evidence to indicate that the operative mechanism driving UFP-mediated vascular dysfunction is not XOR-derived O_2_^●-^ but rather alternative effects of XOR binding and sequestration on the endothelium.

## Material and Methods

### Rodent models

Male and Female, 10-12 week-old, Xdh*^floxed/floxed^*Alb-1cre*^+/-^*(XOR^Hep-/-^) and littermate Xdh Xdh*^floxed/floxed^*Alb-1cre*^-/-^*(XOR^fl/fl^) generated as described previously^37^, along with C57Bl/6J mice (Jackson Laboratory Bar Harbor, Maine; stock no. 000664) and male Sprague Dawley Rats age 10-12 weeks-old (Hilltop laboratories, Scottdale, PA) were used in the study. All animals were maintained in social housing in an American Association for Accreditation of Laboratory Animal Care (AAALAC) approved facility at West Virginia University (WVU) under temperature and humidity regulation on a 12:12 h light-dark cycle. Animals randomly assigned to sham and nTiO_2_ exposure groups and acclimatized to the WVU iTOX facility for 48-72 h prior to starting the experiment with ad libitum access to food and water. Additionally, a small number of mice were further randomized to receive febuxostat or nitrite in their drinking water prior to exposures. XOR^Hep-/-^ and llittermate controls were randomized to receive tail vein injections of purified xanthine oxidase or vehicle. All animal studies were approved by the WVU Animal Care and Use Committee (IACUC).

### Febuxostat treatment (Supp Fig. 1)

After randomization water containing 50mg/L of febuxostat was substituted for normal drinking water. The mice had ad libitum access to the febuxostat water for 5 days prior to a single inhalation exposure along with the access the day of and after exposure. For this experiment water was changed and consumption measured every 2 days.

### Nitrite Treatment (Supp Fig. 2)

After randomization, water containing 50mg/L of NaNO_2_ was substituted for normal drinking water. The mice had *ad libitum* access to the nitrite water for 5 days prior to a single inhalation exposure along with the access the day of and after exposure. Water was changed and consumption measured every day.

### XOR tail-vein injections (Supp Fig. 3)

XOR^Hep-/-^ mice were randomly selected to receive tail vein injections of saline or 5mU of purified XO per estimated ml of blood volume (blood volume= body weight (g) X 0.07), in 50μl of sterile saline (ICU medical, San Clemente, CA) immediately preceding each inhalation exposure.

### Ultrafine Particle Exposure: nano-Titanium Dioxide (Supp Fig. 4)

The nTiO_2_ powder is a mixture composed of anatase (80%) and rutile (20%) TiO_2_, (Evonik, Aeroxide TiO_2_, Parsippany, NJ) with previously defined characteristics: primary particle size (21 nm), the specific surface area (48.08 m/g), and the Zeta potential (−56.6 mV)^44,45^. The nTiO_2_ aerosols were generated by a custom high-pressure acoustical generator (HPAG, IEStechno, Morgantown, WV), which is fed through a Venturi pump (JS-60 M, Vaccon, Medway, MA) to deagglomerated particles and then the aerosol/air mix enters the whole-body exposure chamber. Target mass concentration (10 mg/m^3^ mice, 12 mg/m^3^ rats) was monitored in real-time with a personal DataRAM (pDR-1500; Thermo Environmental Instruments Inc., Franklin, MA). Feedback loops within the software automatically adjusted the acoustic energy to maintain a stable mass concentration during the exposure. Gravimetric measurements were conducted on PTFE filters (0.45 µm pore size) concurrently with the DataRAM measurements to obtain a calibration factor. The gravimetric measurements were also conducted during each exposure to calculate the mass concentration measurements reported in the study. Exposure chamber humidity was maintained to housing facility levels (30– 70%). Sham-control animals were exposed to HEPA filtered air only in an identical chamber (never used for UFP exposures) with similar temperature and humidity conditions. Aerosol size distribution was determined in the exposure chamber using a scanning particle mobility sizer (SMPS 3938; TSI Inc., St. Paul, MN) in tandem with an aerodynamic particle sizer (APS 3321; TSI Inc., St. Paul, MN). Inhalation exposures were conducted in 2 paradigms: 1) Exposures of mice and rats were conducted over 3 consecutive days for either 180 (mice) or 360 (rats) min/day with terminal procedures 24h after the last exposure and 2) To test the therapeutic effects of febuxostat and nitrite, mice received supplemented drinking water as indicated above. After 5 days of supplemented water the mice underwent a single 180-minute nTiO_2_ exposure (or Sham control) with terminal experiments 24h after the exposure. Based on: 1) measurements of the mass median aerodynamic diameter (MMAD) of the aerosol, 2) a measured mass concentration and 3) average animal weight, we conducted aerosol deposition modeling (MPPD Software v 2.11, Arlington, VA). This resulted in a nTiO_2_ alveolar lung burden of approximately 82.7 µg in rats and 16μg in mice over the 3 exposures or a single exposure burden of 5.4 μg in mice.

### Euthanasia and Tissue Harvest

At 24 h post nTiO2 or sham exposure, animals were anesthetized with 5% isoflurane. Whole blood was collected via cardiac puncture and plasma supernatant removed and snap frozen. The mesenteric vascular arcade was then removed and placed in cold physiological salt solution (PSS) and 2^nd^ order arterioles, ∼100μm internal diameter, were excised for vascular reactivity measurements (below). Liver tissue from the medial lobe was then collected and snap frozen.

### Pressure myography of isolated mesenteric arteries

Arterioles were cannulated on glass pipette tips (Living Systems Instrumentation, Burlington, Vermont), and secured with nylon monofilaments suture, as previous^6,46^. Vessels were pressurized to 60 mmHg and then superfused with warmed bubbled PSS containing Ca^2+^ (21% O_2_) and equilibrated to assess the development of spontaneous tone and viability. Arterioles that did not develop > 20% tone were discarded. Arteries were treated abluminal with increasing concentrations of: 1) acetylcholine (Ach: log 10^-9^ – 10^-5^ M), 2) sodium-nitroprusside (SNP: log 10^-9^ – 10^-5^ M), 3) phenylephrine (PE: 10-9 – 10-4 M) and angiotensin II (AngII: log 10^-13^ – 10^-8^) changes in internal vessel diameter were measured to assess vascular reactivity. Additionally, some mesenteric arteries were incubated with febuxostat (10 μM; Axon Medchem BV, Netherlands) for 30 minutes prior to ACh dose response. In a subset of sham XOR^fl/fl^ and sham or nTiO_2_ XOR^Hep-/-^ purified XO at 5mU/ml was perfused into the lumen prior to repeated dose responses to ACh and Ang II. After completion of reactivity studies, the PSS in the isolated vessel chamber was replaced with Ca^2+^ -free PSS and the maximum passive internal diameter of each mesenteric artery was recorded to normalize dilatory responses. Spontaneous tone was calculated by the following equation:

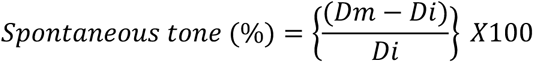

where *Dm* is the maximal diameter and *Di* is the initial steady state diameter recorded prior to the experiment. Active responses to pressure changes were normalized to the maximal diameter using the following formula:

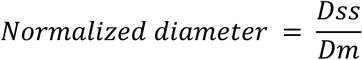

where *Dss* is the steady state diameter recorded during each pressure change. The experimental responses to ACh and SNP are expressed using the following equation:

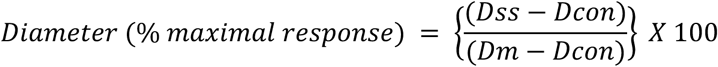

where *Dcon* is the control diameter recorded prior to the dose curve, *Dss* is the steady state diameter at each dose of the curve. The experimental response to PE and IL-33 are expressed using the following equation: where *Dμ* is the unpressurized minimal diameter.

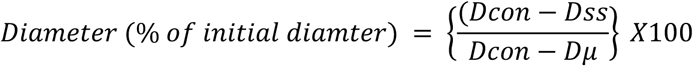

### Intravital microscopy

Rats were anesthetized with Inaction and placed on a heating pad to maintain 37 °C measured by rectal temperature. The airway was kept patent by intubation and the right carotid artery was cannulated to measure arterial pressure. Then an incision on the ventral midline just below the ribs was made to expose the abdominal cavity. Then a 16-20 cm section of the ileum was externalized, small incisions made to flush the section of chyme, and then sutures place to stretch the mesenteric loop over the optical platform. The stage/ optimal platform is continuously superfused with warmed electrolyte solution (119 mM NaCl, 25 mM NaHCO_3_, 6 mM KCl, and 3.6 mM CaCl_2_), supplemented with isoproterenol (10 mg/L) and phenytoin (20 mg/L) to inhibit the peristaltic motion of the intestine and equilibrated with 95% N_2_ and 5% CO_2_. Superfusate was maintained at a flow rate between 4-6 ml/min to minimize atmospheric oxygen equilibration. At this time the animal and platform were transferred to the microscope platform for experimental visualization by our Olympus BX51W1 microscope with Olympus DP71 camera. EDD was measured by iontophoresis delivery of the vasodilatory acetylcholine. The glass micropipettes were filled with 10^-6^ M solution of Ach then advanced to the vessel wall immediately upstream of the visualized area and delivered at 4 different current strengths for 2 minutes with a 2-minute rest period between doses. At the end of all intravital experiments 10^-4^M adenosine was added to the superfusate (final concentration). Digital photomicrographs were captured at steady state response and internal diameter measured by ImageJ and presented as % change in relationship to the maximal adenosine response.

### Xanthine oxidase activity

Xanthine oxidase activity, defined as 1 Unit activity = 1 μmol uric acid generated per min was measured via electrochemical detection of uric acid (UA) following reverse-phase high-performance liquid chromatography (HPLC) as previously described^38,47^. All tissue samples were homogenized in 100 μL PBS containing a protease inhibitor cocktail (Pierce Protease Inhibitor Mini Tablets, ThermoFisher Scientific). Samples were incubated at 37°C for 60 minutes then terminated by addition of ice-cold acetonitrile. The supernatant was carefully removed and transferred into borosilicate glass tubes (14-961-26, Fischer Scientific) and placed in a drying vacuum for 60 minutes. Samples were resuspended in 300 μL of isocratic mobile phase and filtered through a 0.20 μm nylon membrane filter (SLGNX13NK, Fischer Scientific) into plastic snap-top auto-sampler vials (Thermo Scientific). The uric acid content of each sample was measured by electrochemical detection (Vanquish UltiMate 3000 ECD-3000RS) following reverse-phase HPLC separation using a Phenomenex column (Luna 3μm C18(2) 100 A, 150×4.6 mm) and isocratic mobile phase (50mM NaH2PO4, 4mM DTAC, 2.5% methanol, pH 7.0).

### Western blotting

Samples were prepared with laemmli sample buffer and β-mercaptoethanol and run on a 4-20% gradient gel (Biorad, Hercules, California) at 70 V. Samples were then transferred to 0.45 µm nitrocellulose membranes, dried, reconstituted with Tris Buffered Saline (TBS). Membranes were blocked with Li-Cor (Lincoln, Nebraska) TBS blocking buffer, and incubated in primary antibody overnight at 4 °C. Primary antibodies include β-actin (1:1000, Santa Cruz Biotechnology, Dallas, Texas), and xanthine oxidoreductase (1:500, Santa Cruz Biotechnology, Dallas, Texas). Membranes were then washed with TBS containing 1% tween (TBST), incubated in Li-Cor near-infrared secondary antibodies for 1 h at room temperature, washed again with TBST, and finally imaged with the Li-Cor Odyssey Clx. Densitometry analysis was conducted in Image-J (National Institutes of Health, Bethesda, Maryland) reported as target normalized to β-actin and fold change calculated from control.

### Polymerase Chain Reaction

RNA extraction and purification from AML12 cells and liver samples via Qiagen RNA kit and cDNA synthesized by High-capacity RNA-cDNA kit (AB systems). Equal concentration of cDNA was loaded into qPCR reactions in duplicate with 400 nM primer sets and quantified by SYBR green (Bio-RAD). Fold change in ΔΔcT from control was calculated for each sample.

### CUT&Tag sequencing

Experient was conducted with Active Motif® CUT&Tag-IT^TM^ Assay kit (catalog # 53160) following the manufacturers protocol. Human aortic endothelial cells grown to 85% confluency were treated with 5mU/ml XO binding, 5mU/ml glucose oxidase, or vehicle control for 48 hr. Cells were then gently scraped and collected. Roughly 100,000 cells were then bound to concanacalin A beads and incubated with 1μl/ reaction primary antibody against H3K9me^3^ (CUT&Tag validated, Active motif, catalog # 39161) overnight at 4°C with orbital shaking. The next day reactions were incubated with guinea pig anti-rabbit secondary. Then pA-Tn5 Transposases were added to the reaction followed by tagmentation, DNA extraction, and library preparation using the provided i7 and i5 primer combinations for multiplex sequencing. Sequencing was conducted by the Genomic Core facility at Marshal University. Alignment and data analysis were then conducted using Illumina-Basespace software utilizing a FASTQC along with MACs2 and Bowtie for alignment and peak calling.

### Statistics

Data are expressed as either individual points with mean ± SD or as mean ± SE in dose responses in figures while data in the text is represented as mean ± SE. Dose-response curves were analyzed using two-way analysis of variance (ANOVA) with repeated measures and a Holm-Šídák’s multiple comparisons analysis for point-for-point differences when main effect significance was found. All statistical analyses were performed with Graph Pad Prism 9 (San Diego, California). Significance was set at p ≤ 0.05, with n representing the number of vessels, and N representing the number of animals.

## Results

### Impaired Endothelium-Dependent Dilation

As expected, the combination of three inhalation exposures to UFP resulted in significant vasodilatory (Ach) impairment in mesenteric arterioles from both mice (Dose X Group p<0.001) and rats (Dose X Group p<0.001). In the mice dilation was impaired at: 100 nM Ach (28± vs. 15± % dilation, p<0.05), 1 μM Ach (p<0.0001, 44± vs. 22± % dilation), and 10μM Ach (33± vs. 15± % dilation, p<0.001), **Fig. 1A**. In the rats, dilation was impaired at: 100 nM Ach (39± vs. 19± % dilation, p<0.01), 1 μM Ach (60± vs. 28± % dilation, p<0.0001), and 10μM Ach (42± vs. 23± % dilation, p<0.001), **Fig. 1B**. The resting tone that was developed in the isolated vessels from both mice and rats was not significantly altered by UFP exposure, **Fig 1A&B**.

**Fig. 1.**
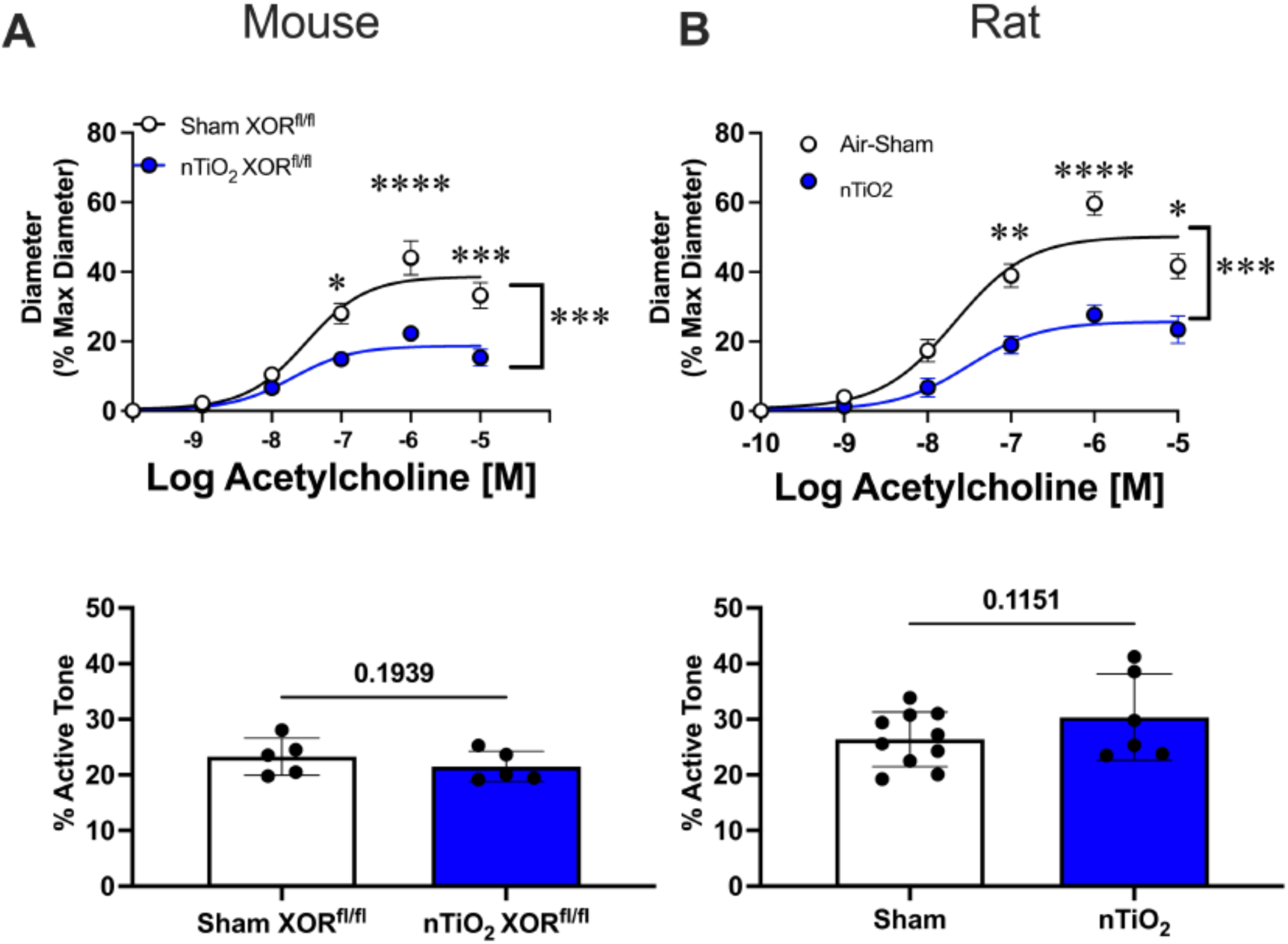
UFPs mediate impairment of endothelial-dependent dilation. **A&B**) Pressurized 2^nd^ order arterioles from sham and nTiO_2_ exposed mice (n=5-6) and rats (n=6-10). Below active resting tone of the pressurized arterioles is shown. * p<0.05, ** p<0.01, *** p<0.001, **** p<0.0001 with,], indicating group X concentration interaction.

### UFP-mediated elevation of circulating XOR

Based on our previous observations that XOR can be upregulated and subsequently released from the liver to the circulation, we assessed XO activity in the plasma and liver of mice and rats following a single exposure to UFP. In mice, UFP increased both plasma (15.7 ± 1.9 vs. 51.6 ± 4.4 μU/mL, p<0.0001 (**Fig. 2A**)) and liver (5.7 ± 0.2 vs. 9.9 ± 0.6 mU/mg tissue, p<0.0001 (**Fig. 2B**)) XOR activity. Likewise, UFP increased XOR activity in plasma (0.3 ± 0.1 vs. 2.4 ± 0.4 mU/mL, p<0.001, (**Fig. 2C**)), and the liver (4.1 ± 0.5 vs. 11.4 ± 1.0 mU/mg tissue, p<0.0001, (**Fig. 2D**)) in the rats. In cultured hepatocytes, 24 h exposure to 0.5 μg/mL of UFP induced a 5-fold increase in XDH gene expression (**Fig. 2E**), a 2-fold increase XOR protein abundance (**Fig. 2F**) and a 3-fold increase in XOR release (4.8 ± 1.6 vs. 15.1 ± 2.8 μU/mL, p<0.05, (**Fig. 2G**)). This coincided with a 2-fold increase in hepatocyte H_2_O_2_ production, **Fig. 2H**. Combined, these findings suggest the UFP inhalation stimulates hepatic upregulation and release of XOR to the circulation.

**Fig. 2.**
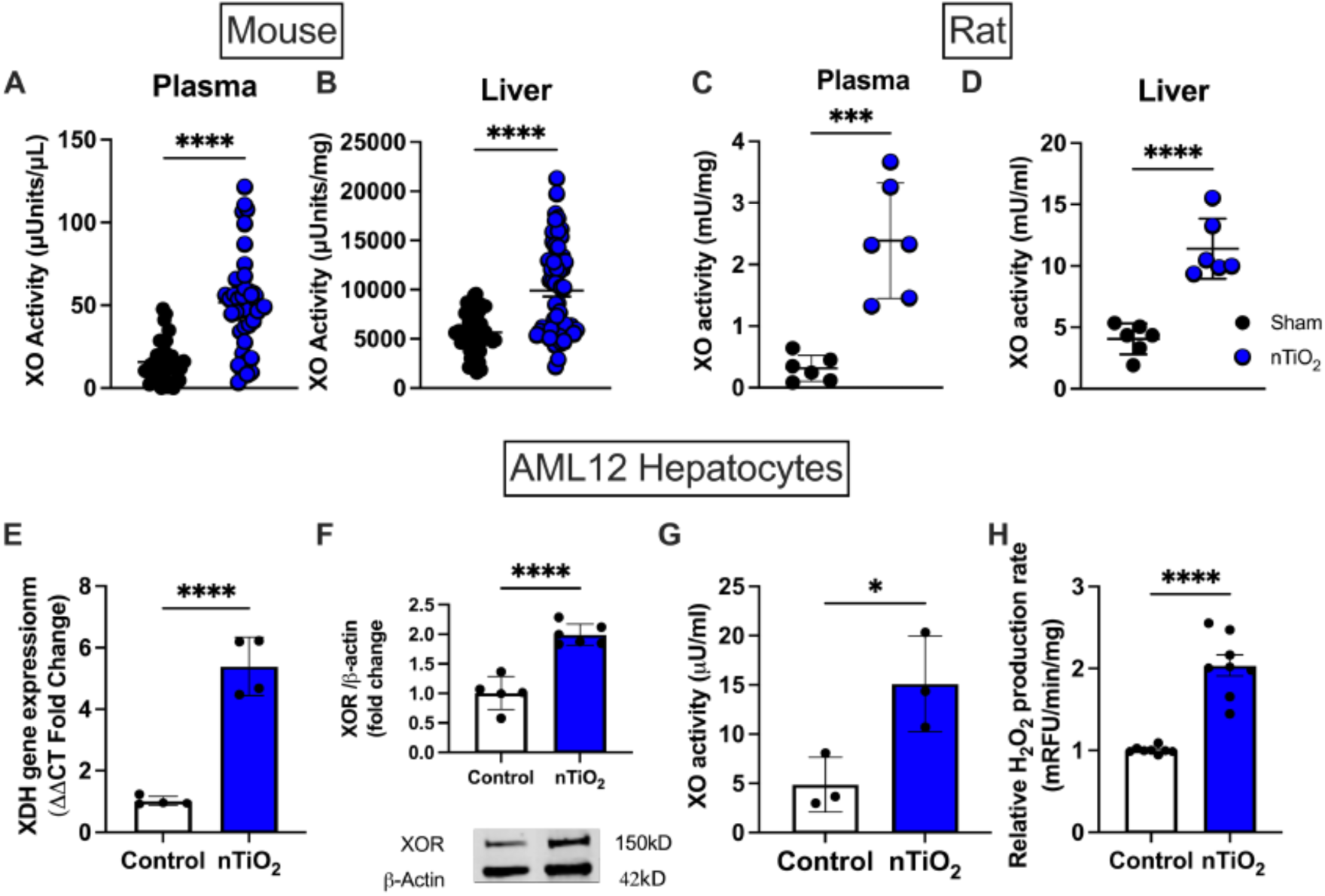
UFPs induced XOR expression and subsequent export from hepatocytes. **A&B**) Activity of XO in plasma and liver from sham and nTiO_2_ exposed mice (n=36-42). **C&D**) Activity of XO in plasma and liver from sham and nTiO_2_ exposed rats (n=6). **E&F**) Gene and protein expression of XOR in cultured AML12 hepatocytes exposed to 0.5 μg/ml of nTiO_2_ for either 6 h (PCR, n=4 in triplicate) or 24 h (WB, n=5-6). **G**) XO release from AML12 hepatocytes measured in the culture media after 24 h (n=3). **H**) H_2_O_2_ production rate measured by CBA following 24 h of 0.5μg/ml of nTiO_2_ (n=8). * p<0.05, *** p<0.001, **** p<0.0001.

### Treatment with either febuxostat or nitrite partial restore vessel dilation in UFP exposed mice

To test therapeutic potential of targeting of XOR in UFP-induced vascular dysfunction we performed chronic treatment with the XOR inhibitor, febuxostat in mice as well as acute administration of febuxostat to rat arterioles *ex vivo*. In mice, UFP impaired vessel responses to Ach was prevented by ad lib administration via the drinking water (max dilation, 20 ± 2 vs. 33 ± 4%, p<0.05 (**Fig. 3A**)), which was expected as febuxostat significantly reduced both liver and plasma XOR activity (liver: 10 ± 0.6 vs. 0.2 ± 0.1 mU/mg, p<0.0001 / plasma: 52 ± 4.4 vs. 6.3 ± 1.7 μU/mL, p<0.0001), **Fig. 3B&C**. Treatment of rat mesenteric arterioles in intravital experiments with 5 μM febuxostat in the superfusate improved dilation (80 nA: 49 ± 3 vs. 25 ± 2 %, p<0.05 / 100 nA: 53 ± 4 vs. 23 ± 1%, p<0.05), **Fig. 3D**. This effect was recapitulated when febuxostat (final concentration of 5 μM) was added to the bath of isolated rat arterioles at the 100 nM Ach dose (27 ± 2 vs. 19 ± 2%, p<0.05) and the 1 μM dose of Ach (37 ± 4 vs. 28 ± 2 %, p<0.05), **Fig. 3E**. Capitalizing on the nitrite reductase activity of XOR, we performed similar experiments by supplementing the drinking water of the mice with 50 mg/L NaNO_2_. Nitrite treatment was equally effective in preventing UFP-induced impairment of vessel reactivity (max dilation: 20 ± 2 vs. 36 ± 5 %, p<0.05), **Fig. 4A**. Surprisingly, nitrite supplementation also reduced XOR activity in the liver and plasma (liver: 10.2 ± 0.6 vs. 3.3 ± 0.1 mU/mg, p<0.0001 / plasma: 52.6 ± 4.4 vs. 24.8 ± 3.8 μU/mL, p<0.0001), **Fig. 4B&C**. Intrigued by this finding, we assessed liver-XOR expression and protein abundance. UFP exposure increased XDH gene expression ∼4-fold (p<0.0001) and XOR protein expression 1.5-fold (p<0.01) which was abrogated by nitrite supplementation, **Fig. 4D&E**. We then treated cultured hepatocytes with 250 μM nitrite, which reduced XDH gene expression by ∼40% compared to vehicle control, **Fig. 4F**. In the aggregate, these data indicate a contributory role for XOR in UFP-induced vascular impairment and that nitrite supplementation diminished this deleterious consequence of particulate exposure potentially by direct interaction with XOR to produce NO or by diminishing the UFP-mediated hepatic upregulation of XDH or both.

**Fig. 3.**
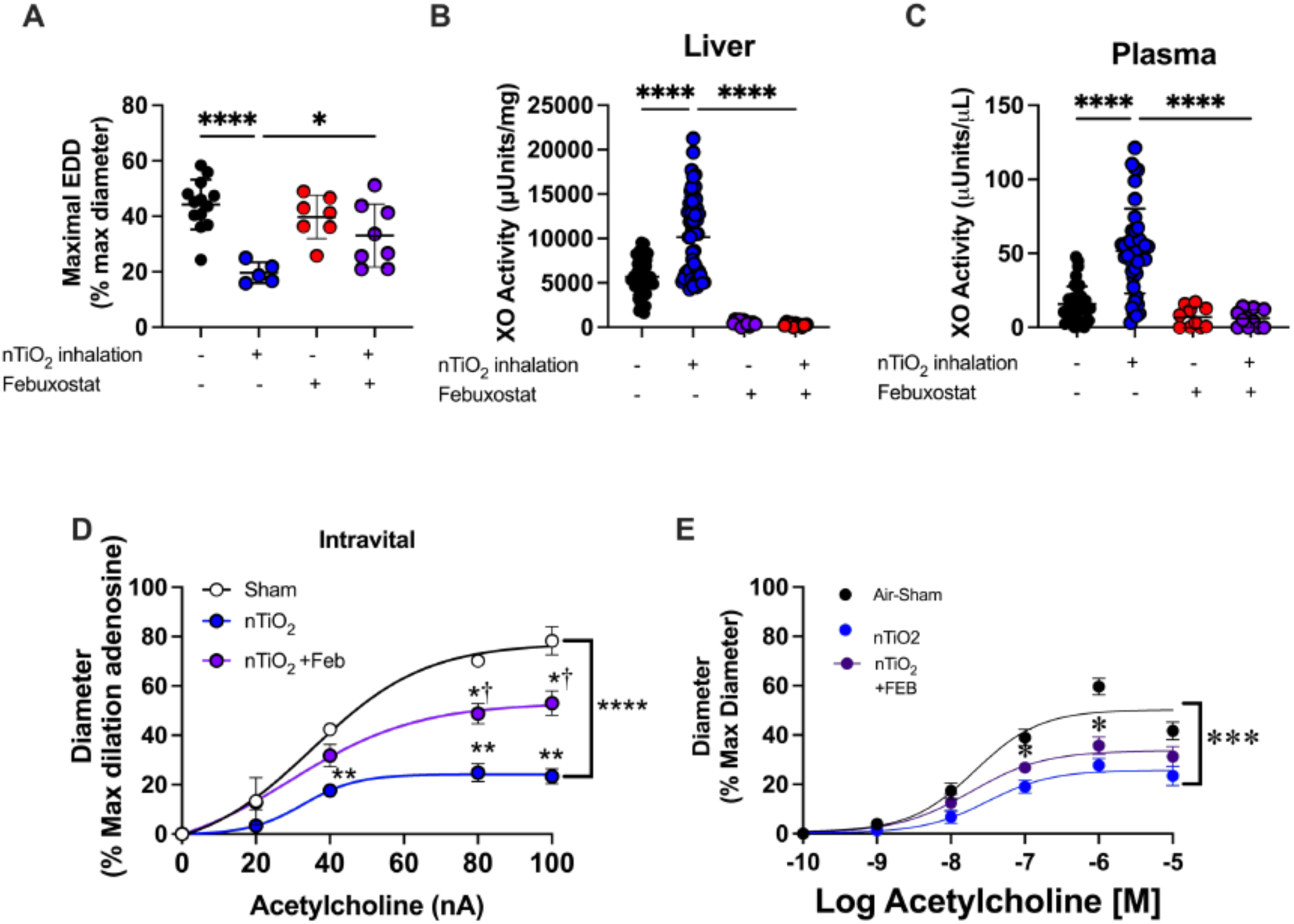
XOR inhibition with febuxostat partially prevents (*in vivo*) or restores (*ex vivo*) UFP-associated vascular impairment. **A**) Maximal dilation to 1μM acetylcholine in pressurized 2^nd^ order arterioles with or without febuxostat administered via drink water (starting 5 days before exposer) in sham and nTiO_2_ single exposed mice (n=5-13). **B&C**) Liver and plasma activity of XO in single exposed mice with or without febuxostat administered via drink water (control as in figure 2 shown again for comparison) (febuxostat treated mice n=11-12). **D**) *In-vivo* intravital assessment of endothelial-dependent dilation of 3^rd^ order mesenteric arterioles. Selected arterioles were stimulated with iontophoresis delivery of acetylcholine then Febuxostat delivered via syringe pump and endothelial-dependent dilation reassessed (Sham+febuxostat had no significant effect and is not depicted on the graph) (n=3). **E**) *Ex-vivo* administration of febuxostat to pressurized 2^nd^ order rat arterioles (n=6-10). * p<0.05, ** p<0.01, *** p<0.001, **** p<0.0001, † p<0.005 compared to nTiO_2_. nA, nano Amps; Feb, febuxostat.

**Fig. 4.**
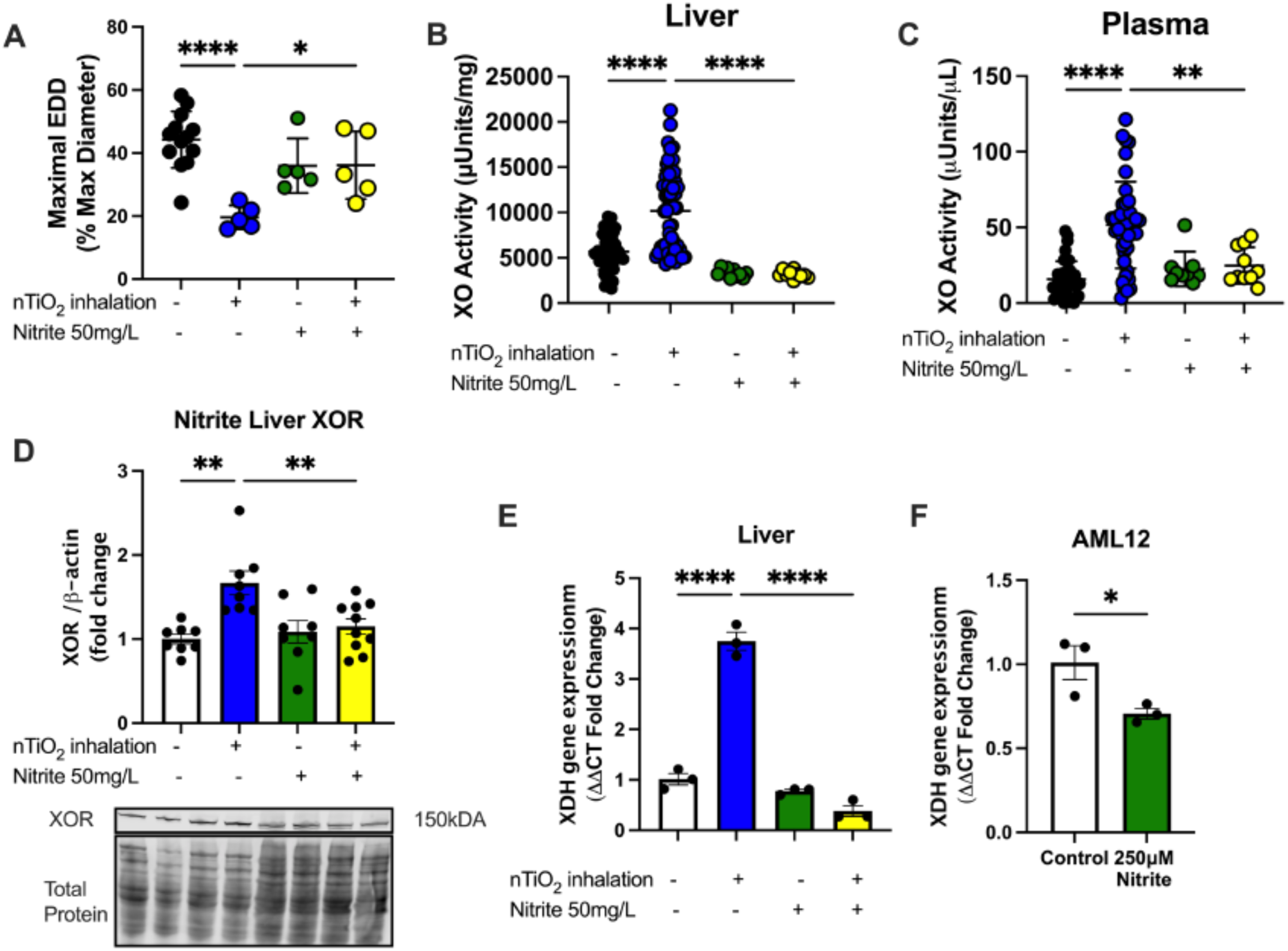
Nitrite supplementation prevents UFP-induced XO upregulation and impairment of endothelial-dependent dilation. **A**) Maximal dilation to 1μM acetylcholine in pressurized 2^nd^ order arterioles with or without nitrite supplementation via drink water (starting 5 days before exposer) in sham and nTiO_2_ single exposed mice (n=5-13). **B&C**) Liver and plasma activity of XO in single exposed mice with or without nitrite supplementation via drink water (control as in figure 2 shown again for comparison) (nitrite treated mice n=9-10). **D&E**) Expression of XOR measured by WB and PCR in liver lysates of single exposed mice with or without nitrite supplementation. **F**) Cultured hepatocytes treated with nitrite for 6 hours and *Xdh* gene expression measured by PCR (n=3 in triplicate). *p<0.05, ** p<0.01, *** p<0.001, **** p<0.0001.

### Hepatic ablation of XOR prevents UFP-induced impairment of arteriole dilation

To examine if hepatic ablation of XOR would impact UFP-induced impairment of vascular function we first confirmed that the resting lumen diameter and wall thickness did not differ significantly between XOR^Hep-/-^ and their littermate controls (XOR^flox/flox^). Surprisingly and unlike the XOR^flox/flox^ littermates, UFP treated XOR^Hep-/-^ mice demonstrated arterioles with greater resting tone (22% ± 2 vs. 29% ± 1, p<0.05) which was not fully reversed by the *i.v.* administration of XOR (29 ± 1% vs. 24 ± 2%, p=0.12) (**Supplement Fig. 5**). Arterioles from Sham-XOR^Hep-/-^ mice exhibited similar vasodilation to Sham-XOR^flox/flox^ controls (max dilation: 44% ± 5 vs. 42% ± 3); however, UFP treated XOR^Hep-/-^ were partially protected exhibiting greater dilation compared to UFP-exposed XOR^flox/flox^ controls at 100 nM Ach (23% ± 2 vs. 15% ± 1, p<0.05) and 1 μM Ach (33% ± 2 vs. 22% ± 1, p<0.05), **Fig. 5A**. Interestingly, re-administration of circulating XO (*i.v.*, 5 mU/mL recombinant XO) in the UFP-XOR^Hep-/-^ reestablished the detrimental effects of UFP. Impaired dilation to Ach was observed at 100 nM (11% ± 2 vs. 23% ± 2, p<0.05), 1 μM (20% ± 3 vs. 33% ± 2, p<0.05), and 10 μM Ach (11% ± 3 vs. 24% ± 3, p<0.05) in the *i.v.* treated XOR group compared to UFP-XOR^Hep-/-^, **Fig. 5B**. Dose response to the endothelium-independent dilator SNP was similar across all groups, **Fig. 5C**. These results suggest that circulating XOR is a key contributor to endothelial-dependent dilation impairment following UFP exposure and does not alter vascular smooth muscle response to nitric oxide.

**Fig. 5.**
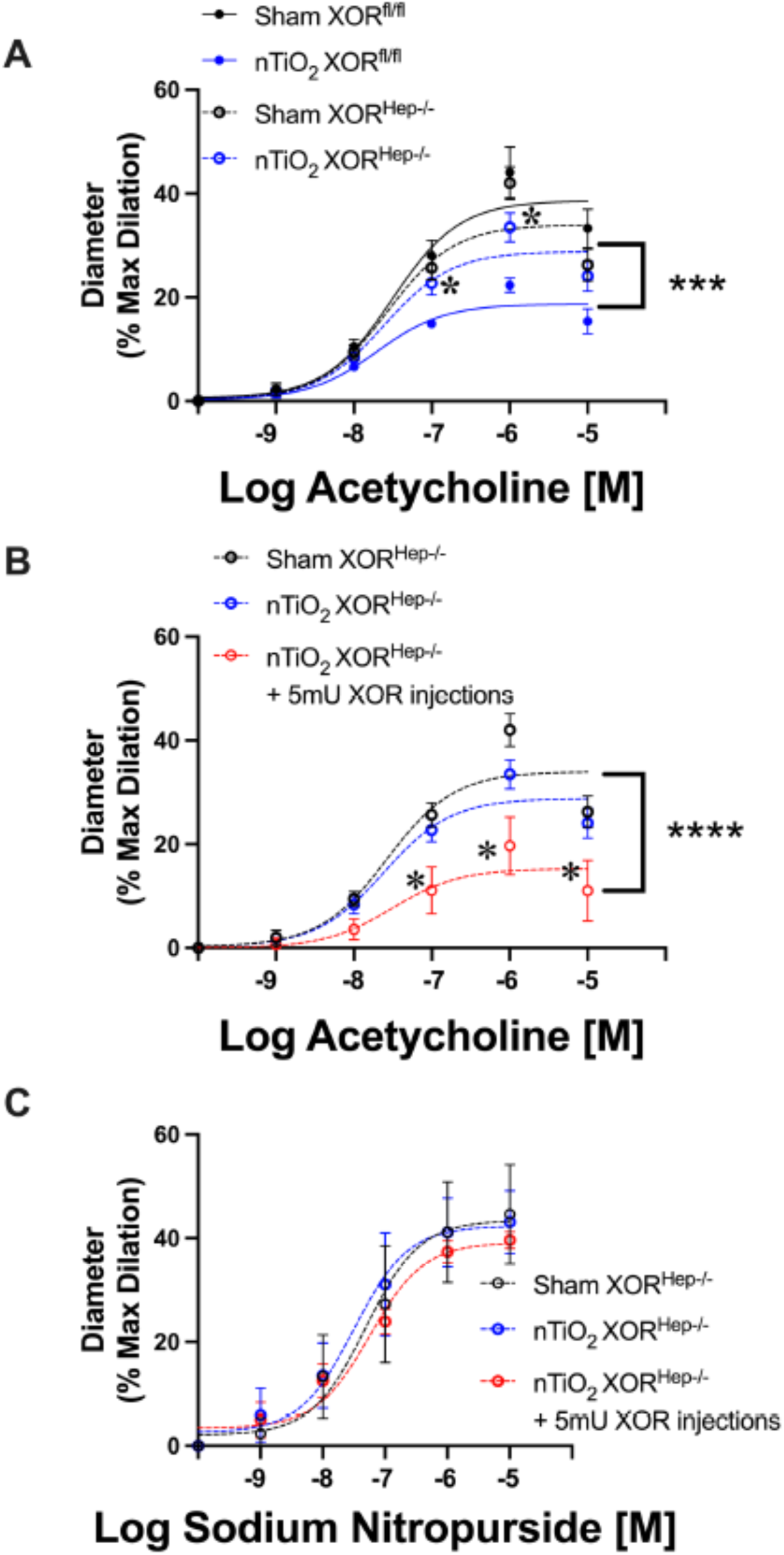
Liver-derived XOR contributes to UFP-associated vessel dysfunction. **A**) Endothelial-dependent dilation in pressurized 2^nd^ order arterioles from sham and nTiO_2_ exposed (3), XOR^fl/fl^ and XOR^Hep-/-^ mice (n=5-6). **B**) Endothelial-dependent dilation in pressurized 2^nd^ order arterioles of XOR^Hep-/-^ mice that had circulating XO reintroduced by iv injections immediately preceding each exposure (n=5). **C**) Endothelial-independent dilation in pressurized 2^nd^ order arterioles from sham and nTiO_2_ exposed (3), XOR^fl/fl^ and XOR^Hep-/-^ mice. In A) * p<0.05 compared to nTiO_2_ XOR^fl/fl^, in C) * p<0.05 compared to nTiO_2_ XOR^Hep-/-^,], indicates 2-way ANOVA group X concentration analysis.

### Acute intralumenal delivery of XOR does not affect endothelial-dependent dilation

To further delineate the potential role of endothelial bound XOR in mediating endothelial dysfunction via diminution of nitric oxide, we delivered purified XOR + 10 µM hypoxanthine to the lumen of arterioles in an *ex vivo* experiment. In healthy arterioles from wild-type littermates, luminal XOR had no effect on Ach dilation (max dilation: 37 ± 7% vs. 44 ± 5%, p =0.43) (**Fig. 6A**). A similar result was observed in both sham (max dilation: 34 ± 4% vs. 42 ± 3%, p=0.36) (**Fig. 6B**) and UFP-exposed (max dilation 30 ± 4% vs. 33 ± 2%, p=0.92) (**Fig. 6C**) XOR^Hep-/-^ mice. Combined, these data suggest luminal XOR does not acutely impair the vasodilatory response to Ach.

**Fig. 6.**
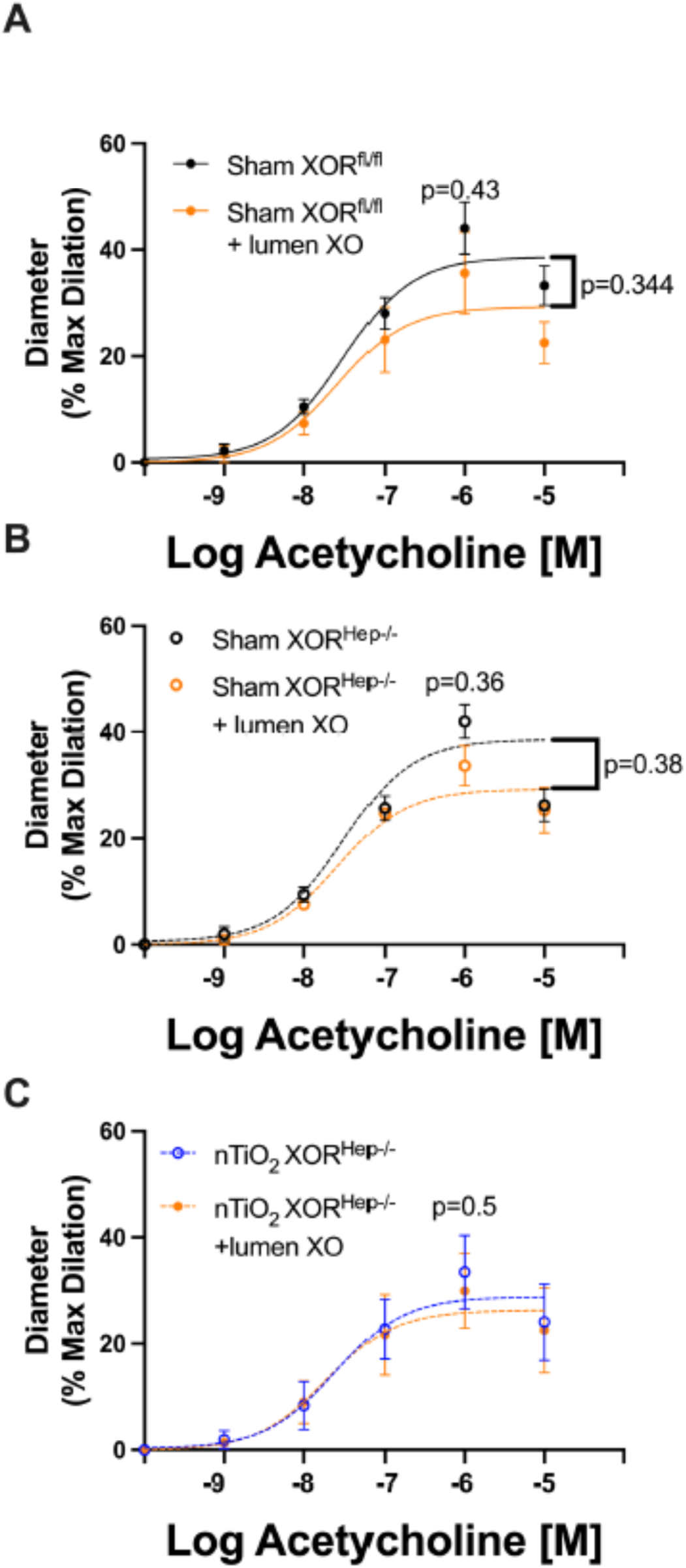
Acute, luminal delivery of XO (*ex vivo*) does not alter endothelial-dependent dilation. Endothelial-dependent dilation in pressurized 2^nd^ order arterioles pre and post lumen delivery of 5mU/ml of XO + 10μM hypoxanthine in **A**) Sham XOR^fl/fl^, **B**) Sham XOR^Hep-/-^, and **C**) nTiO_2_ XOR^Hep-/-^ mice.], indicates 2-way ANOVA group X concentration analysis

### Circulating XO contributes to augmented angiotensin II vasoconstriction

To determine if circulating XOR contributes to vasoconstriction, we performed angiotensin II response experiments. We chose angiotensin II-mediated vasoconstriction as this process is reported to be linked to oxidant production^48^. UFP exposure significantly increased the constrictive response to angiotensin II in control mice (XOR^flox/flox^) (initial diameter: 94 ± 1% vs. 86 ± 1%, p<0.05 (1 nM AngII)) (**Fig. 7A)** whereas there was no significant difference detected in the XOR^Hep-/-^ mice, **Fig. 7B**. Similar to our dilation experiments, restoration of circulating XOR (*i.v.* injection) prior to exposure significantly augmented the angiotensin II response compared to UFP exposed XOR^Hep-/-^ mice (77 ± 4% vs. 94 ± 2%, p<0.05), **Fig. 7C**. Repeating the angiotensin II response following ex-vivo luminal delivery of XO and hypoxanthine (10 μM) did not affect vasoconstriction in sham-XOR^flox/flox^ controls or sham-XOR^Hep-/-^ mice, **Fig. 7D&E**. A modest Group X Dose difference was observed in the UFP exposed, XOR^Hep-/-^ group that were administered exogeneous XOR (p<0.05); however, no difference was detected at any given dose by post-hoc analysis (max response: 94 ± 1% vs. 89 ± 2%, p=0.4), **Fig. 7F**. Taken together, these results suggest circulating XO contributes to UFP-induced augmented angiotensin II responses; however, this contribution is likely not due to the acute presence of the enzyme in the circulation.

**Fig. 7.**
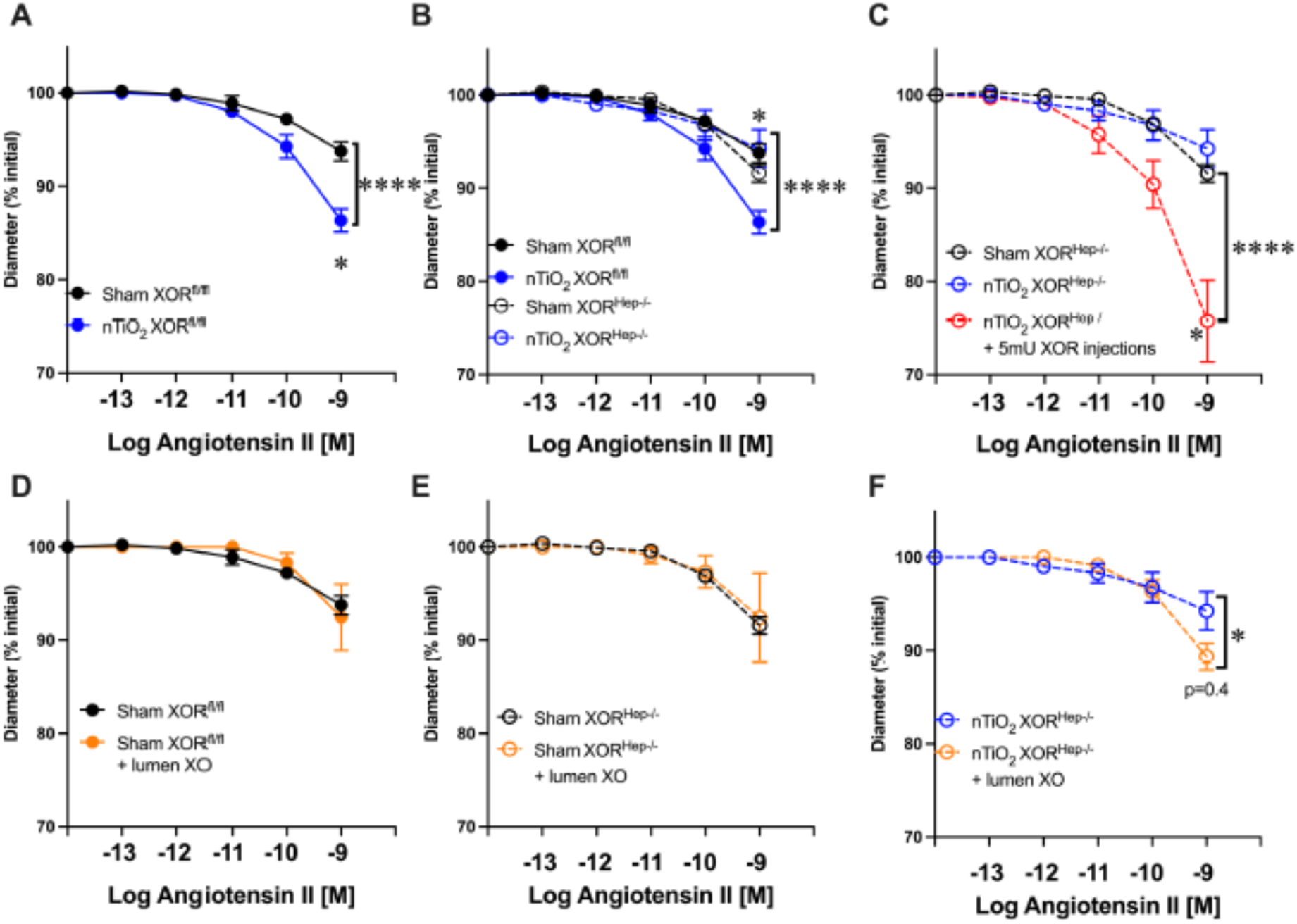
Hepatic-derived XO augments response to angiotensin II. Angiotensin II response in pressurized 2^nd^ order arterioles from sham and nTiO_2_ exposed (3), **A**) XOR^fl/fl^ mice and **B**) XOR^Hep-/-^ mice (n=5-6). **C**) Angiotensin II response in pressurized 2^nd^ order arterioles of XOR^Hep-/-^ mice that had circulating XO reintroduced by iv injections immediately preceding each exposure (n=5). Angiotensin II response in pressurized 2^nd^ order arterioles pre and post lumen delivery of 5mU/ml of XO + 10μM hypoxanthine in **D**) Sham XOR^fl/fl^, **E**) Sham XOR^Hep-/-^, and **F**) nTiO_2_ XOR^Hep-/-^ mice. For (A) *p<0.05 compared to Sham, for (B) *p<0.05 compared to nTiO_2_ XOR^fl/fl^, and for (C) * p<0.05 compared to nTiO_2_ XOR^Hep-/-^.] indicates 2-way ANOVA group X concentration analysis

### Endothelial-bound XOR mediates upregulation of self as well as NADPH oxidase

As our results above indicate that contributions of XOR to UFP-mediated vascular dysfunction is not likely due to the acute presence of elevated enzyme in the vascular compartment and subsequent diminution of NO via O_2_^●-^ production (O_2_^●-^ + ^●^NO → O=NOO^-^), we endeavored to determine potential effects of chronic endothelial glycosaminoglycan (GAG)-bound XOR on oxidant (H_2_O_2_) generation and gene expression (XDH and Nox1). To accomplish this purified XOR was added to the culture medium of human aortic endothelial cells (HAEC) for 1 h to allow for XOR-GAG interaction and then washed to remove all non-bound enzyme; designated as EC-XOR cells. Then the cells were cultured (+/- GAG-bound XOR) with substrate for 24 h. We first analyzed H_2_O_2_ production an found a robust overproduction in the EC-XO which was substantively abrogated by a gentle wash with heparin to compete with and mobilize the GAG-bound XOR as reported previously^49^ and confirms that the H_2_O_2_ was being generated by GAG-bound XOR, **Fig. 8A**. However, even after heparin wash, H_2_O_2_ production was still elevated (∼9-fold) compared to controls. Interestingly, when the heparin washed cells were treated with the XOR inhibitor febuxostat there was a significant diminution (∼60%) in H_2_O_2_ production compared to the heparin washed cells alone; however, overall H_2_O_2_ production was still significantly elevated above the vehicle treated cells, **Fig. 8A**. Analysis of XDH and Nox1 gene expression in the EC-XOR group revealed that both were elevated, **Fig. 8B&C**. In addition, the EC-XOR group also demonstrated increased expression of endothelial nitric oxide synthase, **Fig.8D**. To further explore potential novel avenues by which EC-XOR influences endothelial gene expression we screened for various histone 3 modifications and found a modest but significant reduction in global trimethylation of lysine 9 on histone 3 (H3K9me^3^), **Fig. 8E**. When combined, these results suggest that the presence of elevated endothelial cell-bound XOR mediates alterations in gene expression resulting in the upregulation of self as well as an additional oxidant source, Nox1 and may affect epigenetic pathways/chromatin structure that coalesce to impair vascular reactivity.

**Fig. 8.**
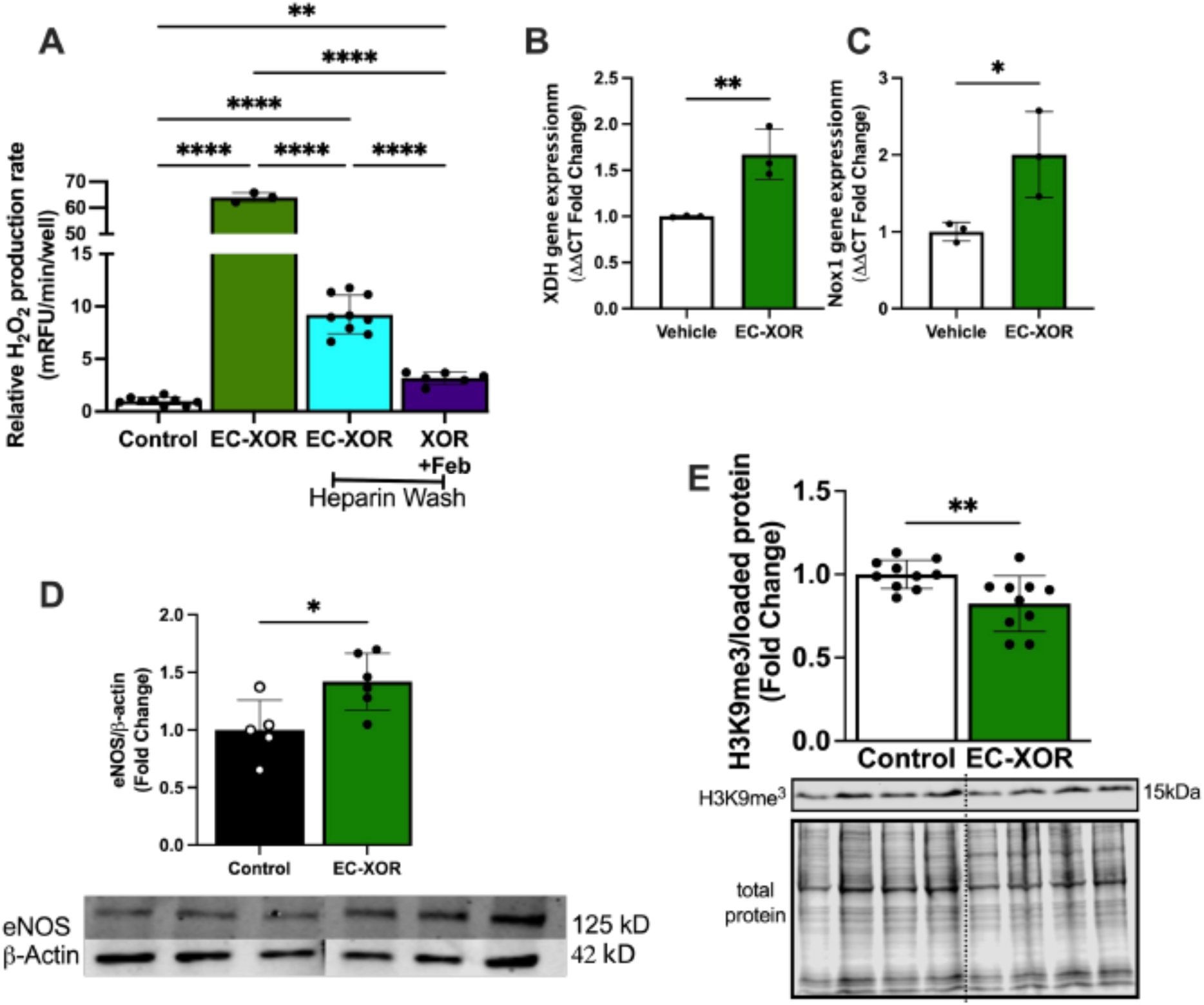
Endothelial bound XO induces *Xdh* and *Nox1* expression and alters histone methylation. Human aortic endothelial cells between passage 3-7 underwent XO binding for 1h then washed to remove unbound enzyme then cultured for 24h. **A**) After 24h H_2_O_2_ production was measured by CBA +/- heparin wash and +/- febuxostat treatment. **B&C**) After 24 h treatment gene expression for *Xdh and Nox1* were analyzed in HAEC. **D**) EC-XOR treatment effect on eNOS expression in HAEC. Finally, western blot analysis for trimethylation of lysine 9 in HAEC 24 h after EC-XO treatment. *p<0.05, ** p<0.01, **** p<0.0001

To test the potential for EC-bound XO as well as the extracellular production of H_2_O_2_ to regulate H3K9me^3^ in endothelial cells we performed CUT&Tag sequencing. HAECs treated with vehicle or 5 mU/ml of GOx or 5 mU/ml XO for 48 hours and processed with the ActiveMotif CUT&Tag assay kit and CUT&Tag validated H3K9me^3^ antibody. Corroborating our western blot findings, we see the H3K9me^3^-associated genome is globally reduced and altered following both treatment with XO and GOx (**Fig. 9A-D**). Analysis of H3K9me^3^ peaks demonstrate a common loss at over 3,500 loci when treating with XO or GOx compared to the vehicle control. 172 peaks were found in common across all 3 treatments. Of interest H3K9me^3^ responses were not identical in the GOx compared to the XO treatments with XO treated HAEC retaining 449 loci (in common with vehicle) that were lost with GOx treatment while GOx retained 29 loci (in common with vehicle) that were lost with XO treatment. In addition to the global loss, unique loci of H3K9me^3^ were discovered following treatment that were not present in the vehicle treated HAEC. GOx and XO treatments resulted in the addition of 16 common H3K9me^3^ peaks, with 20 unique peaks discovered for GOx and 130 unique peaks discovery for XO (**Fig 9E**). Representative IGV plots show the loss of the repressive H3K9me^3^ near the TSS on hallmark genes of inflammation and oxidative stress, ICAM, IL-1β, IL-6, and Nox1 (**Fig. 9F-I**)

**Fig. 9.**
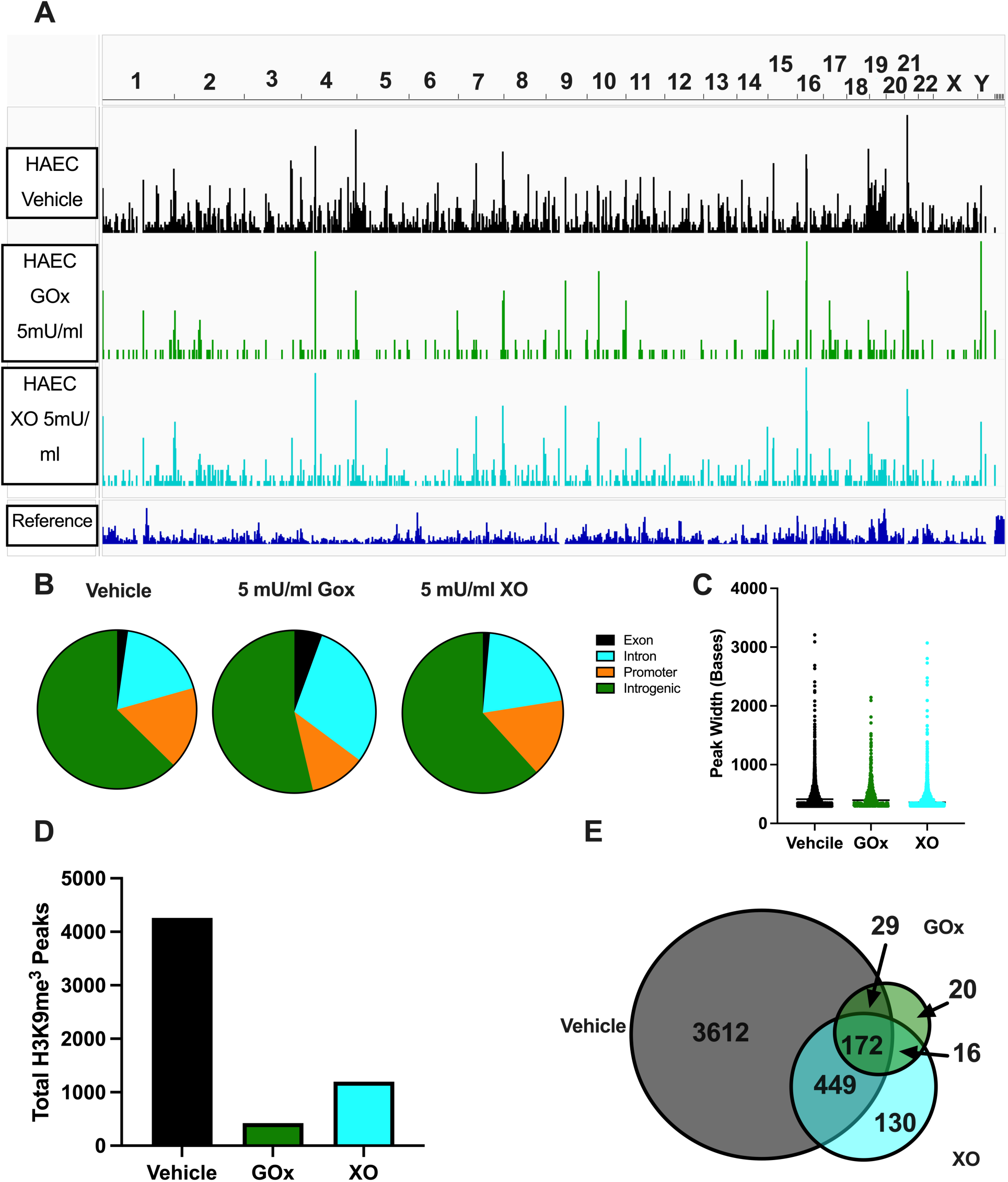
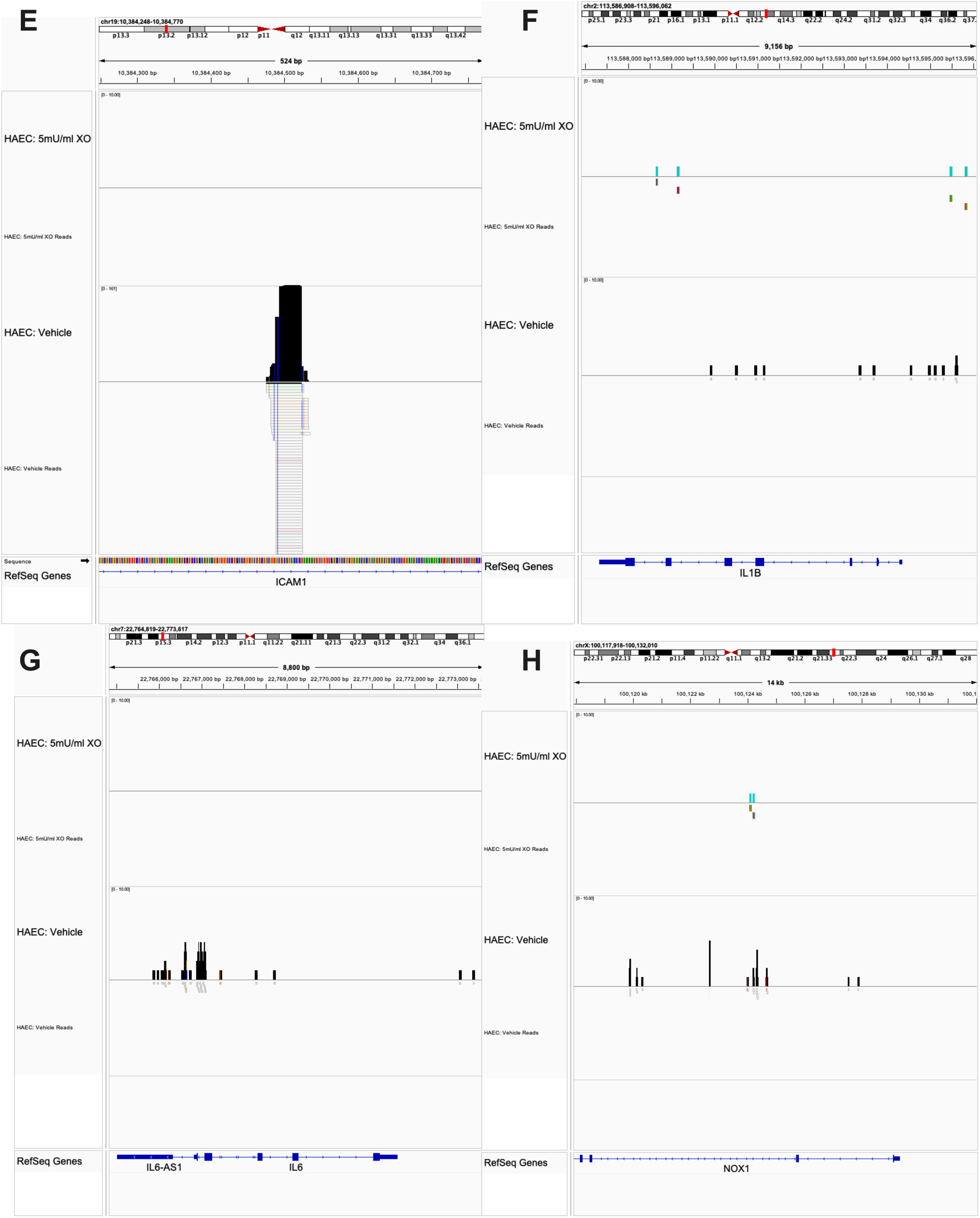
H3k9me^3^ CUT&Tag in human aortic endothelial cells. **A**) Combined peak tracks for vehicle, 5mU/ml GOx, and 5mU/ml of XO depicted in the Integrative Genome Visualizer (IGV). **B-D)** Relative genomic sight and peak width of H3K9me^3^ peaks. **D-E**) Total number of called peaks in the combined data set with differential peak identification venn diagram with overlapping area represents shared peaks while nonoverlapping areas represent peaks unique to that treatment. **F-I**) Representative peaks showing loss of H3K9me^3^ on ICAM, IL-1β, IL-6, and Nox1, IGV. n=4 per with >40 million aligned reads (>98% alignment)

## Discussion

Epidemiologic data strongly support a link between inhalation of UFP and cardiovascular pathologies^3,4^. The chemical and structural composition of commonly encountered UFP in the environment is heterogenous and varies daily which reveals considerable issues with real-time, real-life exposure modeling in preclinical models. To address this issue, our group has utilized nTiO_2_ as a surrogate to produce consistent and reliable UFP exposures^5,6,11,46^. Further substantiating our rationale for using nTiO_2_ is its use in an ever-growing list of industrial, consumer, and medical products which enhances exposure both occupationally and via product handling/consumption. Our group has described systemic oxidant-related vascular impairment linked to nTiO_2_; however, the source and mechanisms of oxidant-induced vascular impairment remain underdeveloped^6,10,50^. With this as a backdrop, we posit that XOR may be a critical contributor of the enhanced oxidant load reported in the above referenced reports. We propose this as XOR has been tightly linked with in cardiovascular pathologies, its presence in the circulation, and our recent studies demonstrate that it can be upregulated and released from hepatocytes upon inflammatory stimulation^38,47^. In addition, we and others have reported that once released to the circulation, XDH is rapidly converted to XO and this XO can avidly (*K_i_* = 6 nM) bind to the endothelial glycocalyx; specifically, to heparin-containing glycoproteins via a heparin binding sites on the surface of the enzyme^51,52^. This binding to and sequestration of XO on the apical surface of the endothelium creates a local milieu that is primed for production of oxidants that can participate in loss of vascular homeostasis.

To test the hypothesis that XOR contributes to UFP-associated vascular dysfunction, we first validated our previous nTiO_2_ studies, using rats, in a mouse model to establish a similar vascular-impairment phenotype. Results from these experiments revealed that exposing mice to nTiO_2_ for 3 h (total alveolar lung burden of 5.4 μg/exposure) produced a similar endothelial impairment to rats exposed to 27.3 μg/exposure. Having validated our vascular phenotype, we interrogated the impact of exposure on hepatic and circulating XO levels. While associations between toxicants and XOR levels have been reported, these observations are limited both in scope and number. For example, heavy metal exposure in both humans and fish is linked with elevation of XOR and UA levels^41–43^ and compounds from cigarette smoke induce XOR upregulation^53^. Our findings support these previous observations and provide initial evidence that inhaled particles elicit signals in hepatocytes to upregulate XOR (**Fig. 2**). How a primary insult in the lung is translated or communicated to the liver is an interesting question that warrants deeper investigation; however, evidence exists suggesting inhaled UFP particles can reach the circulation and have a propensity for hepatic accumulation^39,40,54^. When combined, these previous observations and our data in cultured hepatocytes (**Fig. 2F-H**) support a potential direct stimulation and release of XOR by nTiO_2_. If this process is operable *in vivo* and/or requires the contributions of previously defined redox-signaling pathways^38^ is yet to be determined.

Nitrite/nitrate supplementation has been used to address a wide array of vascular inflammatory disease processes by elevating the local NO levels as a consequence of reductive processes that act to directly convert NO_3_^-^ and NO_2_^-^ to NO^55^. A key source of this reductive power for converting NO_3_^-^ and NO_2_^-^ to NO is XOR where electrons provided by either hypo/xanthine or NADH can be used to reduce NO_3_^-^ and NO_2_^-^ at the molybdenum cofactor (Mo-co)^55–58^. This nitrite reductase functionality of XOR is operative under inflammatory conditions that limit the canonical, eNOS, path to NO generation while also diminishing XOR-derived oxidant generation due to electrons being shuttled to nitrite and not O_2_^59^. For example, nitrite supplementation has been effective in providing improvement in models of vascular disease^60,61^ while concomitant inhibition of XOR has abrogated this effect thus demonstrating XOR-dependent contributions to the salutary actions (NO generation) of nitrite supplementation^62–65^. As such, we hypothesized that, in the context of UFP-mediated vascular dysfunction, we could capitalize on the elevated abundance of XOR combined with its nitrite reductase activity by elevating nitrite levels and switching XOR product identity from oxidants to beneficial NO.

When mice were subjected to nitrite supplementation prior to UFP exposure, vessel dysfunction (impaired EDD) was significantly improved, **Fig. 4**. We also observed that nitrite supplementation resulted in diminution of XOR activity in both the plasma and liver and we surmised this was either due to nitric oxide-mediated inactivation of the enzyme^66^ or transcriptional down regulation. When we interrogated the latter in the livers of nitrite-treated mice as well as in nitrite-treated, cultured hepatocytes, we observed diminished XDH mRNA levels, **Fig. XX**. This could mean one of two things: 1) nitrite mediates a repression of *Xdh* gene transcription or 2) nitrite supplementation mediates a reduction in XDH mRNA stability. Regardless of the direct cause, the reduction of XDH message and protein levels in the liver was also associated with diminished circulating XO, **Fig. 4**. While this observation of the association between nitrite supplementation and XDH message was not expected, it does not diminish the impact on vessel function but rather sheds new light on alternative contributions that are yet to be fully vetted via additional experimentation.

Our results with the hepatocyte-specific XOR knockouts make it clear that liver-derived XOR is a significant contributor to UFP-induced impairment of vascular function. This is exemplified by the preservation of the EDD response to Ach and a similar Ang II response to that of the sham-littermate controls in the UFP-exposed XOR^Hep-/-^ mice. Gain of function experimentation via direct restoration of circulating XO in our XOR^Hep-/-^ mice, resulted in UFP-impairment of vascular function. These data serve to strengthen our supposition that UFP exposure stimulates hepatocytes to actively release XOR to the circulation where XO subsequently contributes to vascular impairment. This is partially supported by our previous work in a model of experimental heme crisis where either iron or hemin or hemopexin-hemin all triggered hepatocellular upregulation of XDH, XOR protein and export to the circulation^36,38,47^. This same set of experiments also revealed results that challenge long-standing dogma regarding the presence of XO in the vascular compartment and specifically when it is associated with endothelial glycosaminoglycans. For example, in non-hemolysis-related vascular inflammatory processes, elevation of XO has long been relegated to a deleterious role predicated upon its production of superoxide (O_2_^•-^) and then subsequent diffusion-limited reaction with NO which results in diminution in bioavailable NO and ultimately, loss of capacity to dilate. However, our data countervail this concept as acute delivery of exogenous XO plus substrate to the vessel lumen does not alter response to Ach indicating that O_2_^●-^ generated by XO in this environment does not impact endothelial derived NO and its diffusional progress to the smooth muscle, **Fig. 6&7**. Considering that, if indeed the XO-derived, and negatively charged, O_2_^●-^ was generated on the apical surface of the endothelium, then it would be difficult to conceive how it would traverse the plasma membrane and intercept NO as it diffused to the basolateral surface on its way to the smooth muscle. These data (**Fig. 6&7**) and the concept outlined above potentially reveal that the presence of XO on the endothelium may be affecting vascular responses to dilation via a route alternative to direct effects on NO. In addition, in this acute study we delivered XO to the lumen of a canulated vessel in non-oxygenated PSS (not actively aerated with O_2_) whereas previous published reports delivered XO to the superfusate surrounding the vessel or utilized heparin removal of bound XO in aortic ring experiments^35,67^. The superfusate bath for aortic rings or even canulated microvessels is hyperoxygenated (bubbled with 95% or 21% O_2_, respectively) when compared to normal, physiologic environments. In these previous experiments XO free in solution with 21% bubbled O_2_ or XO bound to the endothelium of aortic rings with 95% bubbled O_2_ would produces considerably more O_2_^●-^ than when bound and at lower levels of oxygen tension relevant to the *in vivo* setting where bound XO oxidant production is >80% H_2_O_2_^68^. In a similar vein oxidant production is shown to mediate Angiotensin II-associated vasoconstriction^48^. Our results indicate circulating XO augments the Angiotensin II response, but this too was not observed when XO plus substrate was delivered acutely to the vessel lumen. As our results above we derived from an acute *ex vivo* exposure to XO, we postulate that the absence of an effects in this setting may be rooted in the difference between this model and our *in vivo* model where XO is present in a chronic manner where it can potentially impact endothelial cell function over time.

The process by which chronically elevated XO induces vascular impairment obviously requires further exploration; however, we propose two potential mechanisms of this chronic effect on endothelial function. First, and as previously shown, XO can be taken up by endothelial cells which may take time to accumulate and become spatially oriented with NO pools. While circulating/endothelial bound XO may not contribute acutely, cellular sources of XO may acutely impair EDD. This idea is supported by data herein as well as numerous reports demonstrating acute administration of XO inhibitors improves EDD responses to Ach^35,36,67^. Importantly, it appears both feboxostat and allopurinol acutely improve EDD, which circumstantially supports that endothelial GAG-bound XO doesn’t acutely impair EDD, as allopurinol is not effective at inhibiting GAG-bound XO even at concentrations far above those achievable in the clinic (200-400 μM)^69^. This would suggest that acute pharmaceutical inhibition is having its effects through targeting intracellular pools of XOR. Second, H_2_O_2_ produced by XO on the apical surface of the endothelium, enters the cell via aquaporins^70^, to initiate redox-mediated changes to gene expression/enzymatic processes. Our CUT&Tag data strongly supports that extracellular H_2_O_2_ mediates significant alterations to chromatin structure, **Fig. 8&9**. This is supported by recent evidence that H_2_O_2_ can alter epigenetic gene regulation^71,72^. Our data suggests that EC-bound XO drives redox-mediated loss of H3K9me^3^, which may prime the endothelial cells to upregulate inflammatory genes. These data provide critical insight as to how XO/redox-signaling may initiate chromatin remodeling in the context vascular disease initiation/progression. While the focus of this manuscript was on XO’s redox-mediated it should be noted that EC-bound XO caused unique H3K9me^3^ alterations dependent from those seen with chronic delivery of extracellular H_2_O_2_, suggesting the physical interaction, uric acid production, or depletion of xanthine may play a unique role in endothelial cellular signaling. Our data strongly suggests a link between XO/redox-signaling and H3K9me^3^ in endothelial cells, future work aimed at uncovering the mechanisms linking XO/ redox-signaling to the writing, reading, and erasing of histone methylation should yield exciting results and potentially identify novel therapeutic targets in redox-mediated cardiovascular pathologies.

*In toto*, the result outlined above provide evidence that hepatic-derived, circulating XO contributes to the impairment of vascular function allied to UFP exposure. We believe this process involves chronic elevation of circulating and endothelial-associated XO that imparts alterations in endothelial cells mediating diminution in capacity to dilate. For example, new evidence herein indicates endothelial-bound XO contributes to altered gene expression potentially via alterations in H3K9 methylation. Importantly, nitrite supplementation is an effective therapeutic to address vascular dysfunction where XO is found to be contributory including, but not limited to, inhalation exposure to particulates where potential through downregulation of XOR expression and release from hepatocytes. And finally, we recognize that there are several areas in which further experimentation is requisite to identifying mechanisms underpinning our observations and currently planning next steps.

## Acknowledgements

Funding: NIH U54GM104942 and AHA 23CDA1038976 (E.D.); NIH HL153532, DK124510 and 19TPA34850089 (EEK); NIH ES015022 (TRN); NIH ES031253 (SH); K01 ES015022 (EB)

## Abbreviations

GAGs: glycosaminoglycans
H_2_O_2_: hydrogen peroxide
^●^NO: nitric oxide
NO_2_^-^: nitrite
NO_3_^-^: nitrate
O_2_^●-^: superoxide
XDH: xanthine dehydrogenase
XO: xanthine oxidase
XOR: xanthine oxidoreductase

## Notes

### Competing Interest Statement

The authors have declared no competing interest.

