## Supplemental Figures for "Liver-Derived, Circulating Xanthine Oxidoreductase Drives Vascular Impairment Associated with Inhalation of Ultrafine Particulates"

### Supplemental Figure Legends

**Suppl. Fig. 1. Febuxostat treatment**. **A**) Schematic of febuxostat single exposure experiment. **B**) Measured water consumption.

**Suppl. Fig. 2. Nitrite treatment**. **A**) Schematic of nitrite supplementation single exposure experiment. **B**) Measured water consumption and **C**) plasma $NO_x$ concentration. **** $p<0.0001$

**Suppl. Fig. 3. Three-day UFP exposure with vehicle of 5mU XO *i.v.* delivery**. Schematic of 3-day exposure experiment with i.v. administration of XO.

**Suppl. Fig. 4. Ultrafine particle ($nTiO_2$) characterization**. **A**) Representative tracing of mean aerosol concentration with the black line being the observed concentration and the red line indicating the targeted set-point (10mg/$m^3$, for a mouse exposure). **B**) particle size distribution measured by tandem scanning mobility particle sizer (light grey) and aerodynamic particle sizer (dark grey) count median diameter 116 nm ± geometric standard deviation 2.11, **C**) high-resolution electrical low-pressure impactor, count median diameter 169 nm ± geometric standard deviation 1.94, **D**) along with a micro-orifice uniform deposit impactor with mass median aerodynamic diameter of 1.03mm ± geometric standard deviation 2.57.

**Suppl. Fig. 5. Resting tone of mesenteric isolated 2$^{nd}$ order arterioles**. Isolated 2$^{nd}$ order arterioles hung in living systems pressure myography rig and allowed to develop spontaneous tone.

**Supplemental Figures**

Supplemental Figure 1.

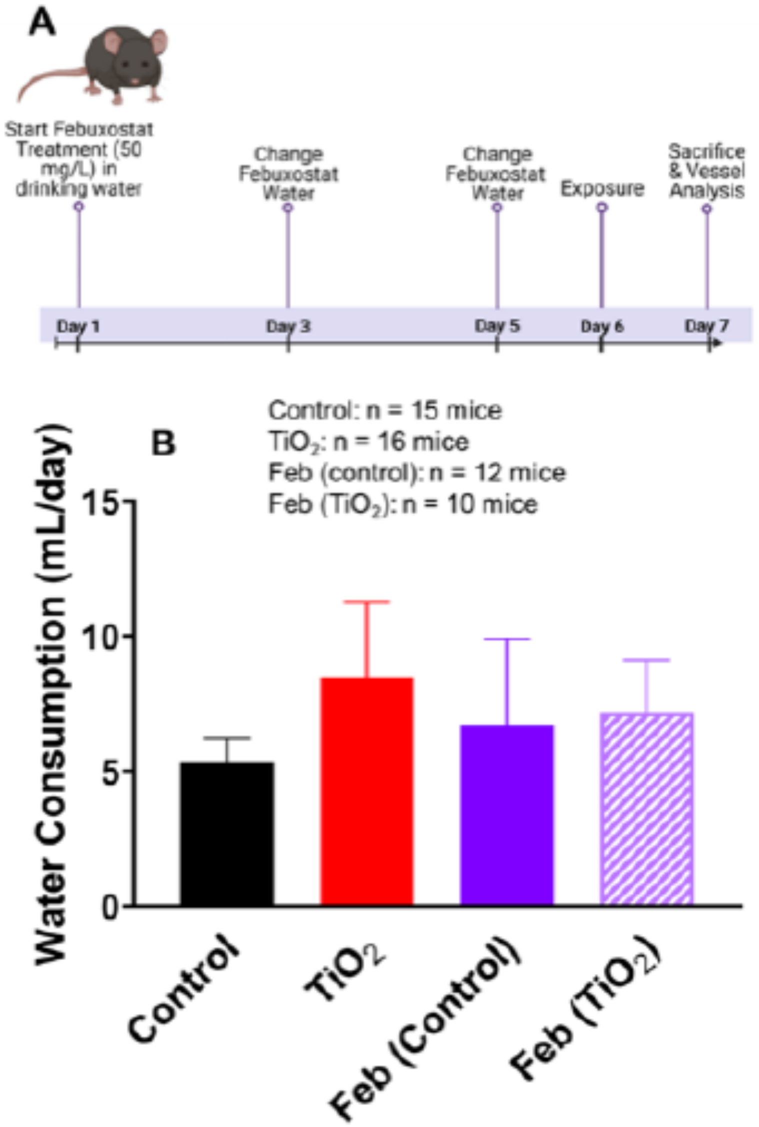

**Supplemental Figure 2.**

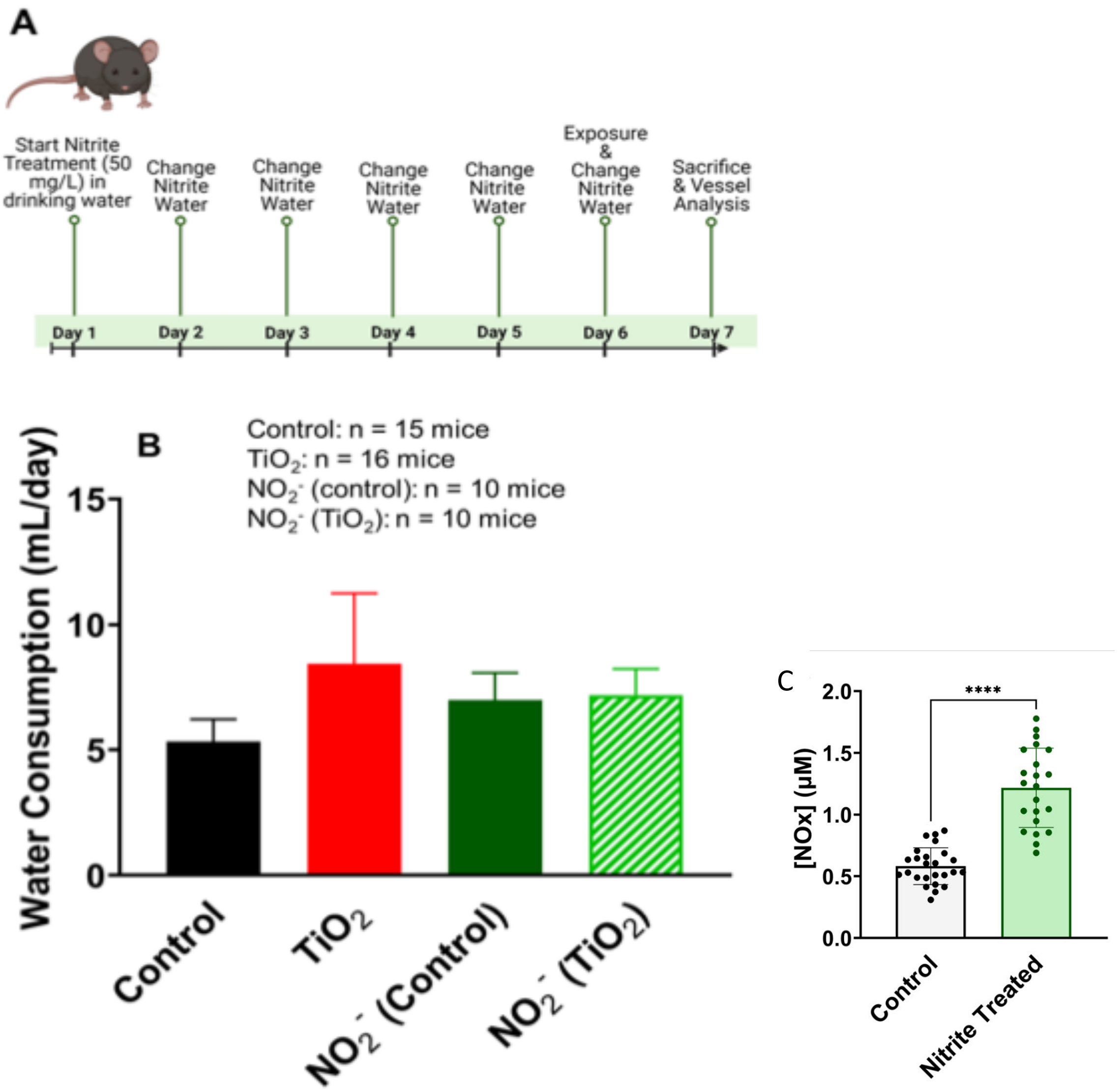

### Supplemental Figure 3.

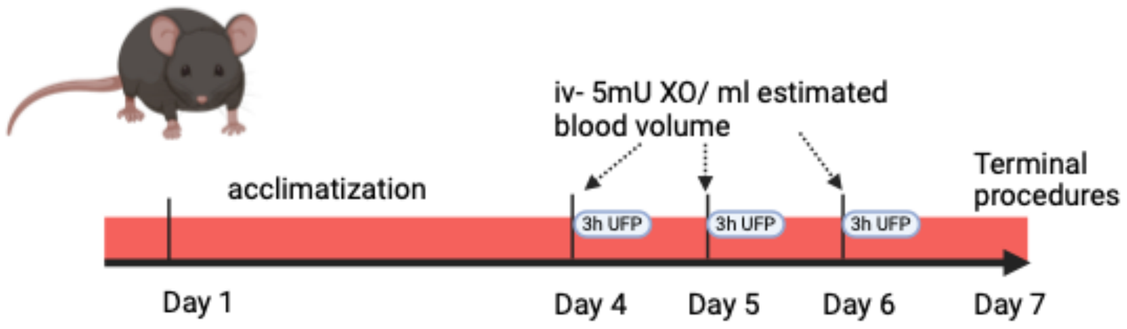

**Supplemental Figure 4.**

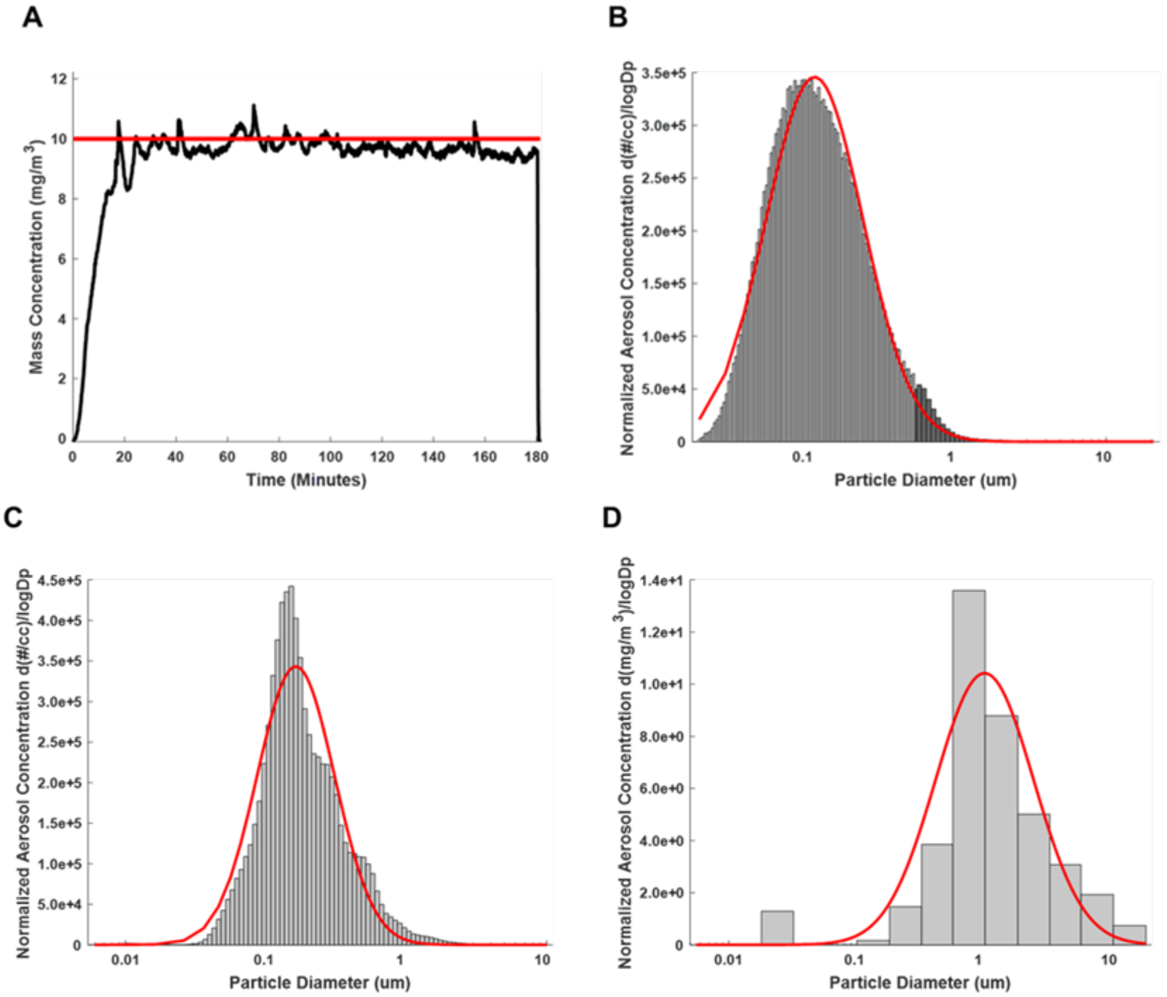

Supplemental Figure 5.

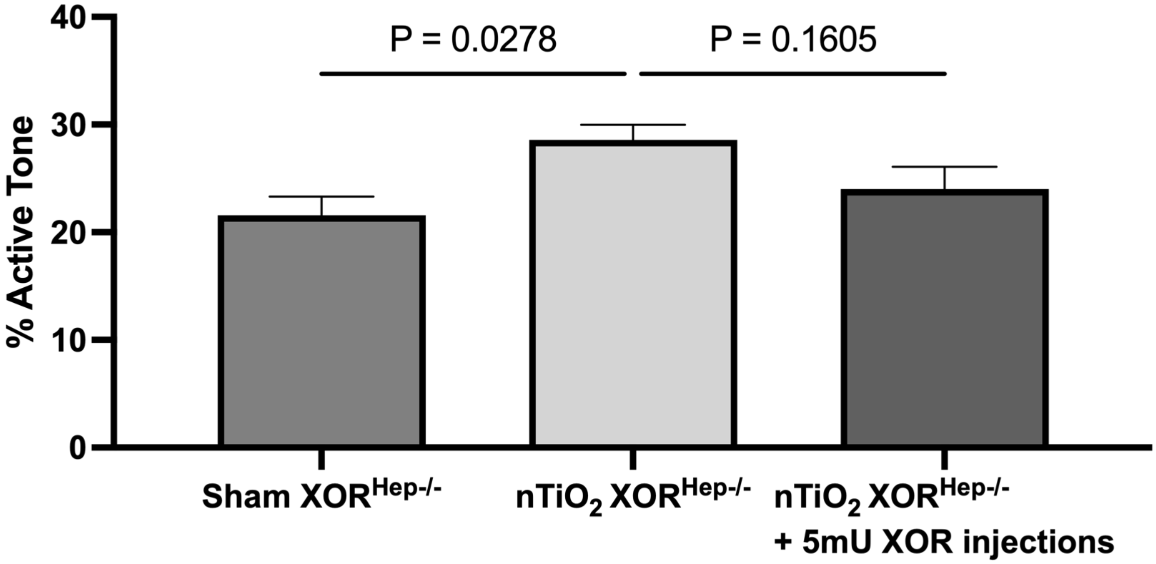
